# The small molecule inhibitor SU056-mediated targeting of YB1 inhibits the Rb pathway in triple negative breast cancer tumors

**DOI:** 10.64898/2026.09.05.749600

**Authors:** Wei Wang, Yeaji Kim, Xiaotong Zhao, Kristen Weber Bonk, Ruth Keri, Bin Su, Khalid Sossey-Alaoui

**Author notes:** **Corresponding Author:** Khalid Sossey-Alaoui, PhD, Associate Professor, Department of Medicine, Case Western Reserve University School of Medicine, Division of Cancer Biology, MetroHealth Medical Center, Rammelkamp Center for Research, R457, 2500 MetroHealth Drive, Cleveland, OH 44109, (lab), https://case.edu/cancer/members/member-directory/khalid-sossey-alaoui.

## Abstract

**Background:** Triple-negative breast cancer (TNBC) tumors lack expression of estrogen receptor (ER), progesterone receptor (PR), and HER2, limiting the availability of targeted therapeutic options. As a result, TNBC is characterized by a high propensity for metastasis, rapid recurrence, and poor overall prognosis. Despite significant research efforts, the molecular mechanisms that drive TNBC progression and metastasis remain incompletely understood.

Y-box binding protein-1 (YB1) is a multifunctional DNA/RNA-binding protein that regulates transcription, translation, and mRNA stability. Aberrant activation of YB1 has been implicated in multiple oncogenic processes, including proliferation, survival, invasion, metastasis, and therapy resistance in several cancer types, including TNBC. Our investigations have revealed a central role for YB1 in regulating key hallmarks of cancer that drive TNBC tumor progression and metastatic dissemination.

**Methods:** Cell viability and drug sensitivity in TNBC cell lines were assessed using the MTT assay to determine IC₅₀ values for pharmacologic inhibitors. Western blotting and quantitative RT-PCR were used to quantify protein and mRNA expression levels in TNBC cell lines and tumor samples. Oncogenic phenotypes were evaluated using 2D colony formation assays, 3D tumorsphere growth assays, limiting dilution, and transwell migration assays to assess proliferative, stem-like, and migratory capacities. Cell cycle progression was analyzed using flow cytometry–based assays. In vivo therapeutic efficacy was evaluated using preclinical xenograft and patient-derived xenograft (PDX) mouse models of TNBC to assess tumor growth and metastatic progression following treatment.

**Results:** Our studies demonstrate that the small-molecule inhibitor SU056, which specifically targets YB1 for degradation, significantly suppresses the oncogenic behavior of TNBC cell lines and tumors. Pharmacologic inhibition of YB1 reduced TNBC cell proliferation, clonogenic growth, tumorsphere formation, stemness and migratory capacity. Importantly, combined treatment with SU056 and the CDK4/6 inhibitor palbociclib enhanced the inhibitory effect on TNBC cell growth and tumor progression compared with either agent alone. Mechanistically, both genetic and pharmacologic targeting of YB1 suppressed TNBC tumor growth and metastatic potential through modulation of the Cyclin D/CDK4/6/Rb signaling axis, leading to inhibition of cell cycle progression and induction of G1 arrest.

**Conclusion:** Collectively, these findings identify SU056/CDK4/6i combination as a potential therapeutic option for the treatment of Rb^+^-TNBC.

## Introduction

Breast cancer (BC) is the most diagnosed cancer in women and is the leading cause of cancer-related death worldwide [1]. To date, no cure has been found for advanced BC, with a mediocre median patient survival of only 1.5 to 3 years. Effective therapies have been further limited by the heterogenic nature of BC tumors, which can be subdivided into several genetically and molecularly distinct subtypes [2–4]. Hormone-receptor positive (ER-α^+^ and PR^+^), which constitute the vast majority (∼70%) have the best prognosis when detected early, thanks to their effective response to targeted hormone and chemo/radiation therapies. However, those classified as triple negative breast cancers (TNBCs) are especially lethal due to their highly metastatic behavior and propensity to recur rapidly [5]. As a group, TNBCs lack expression of hormone receptors, ER-α and PR, and are ErbB2/HER2 [6], which has hindered the development of FDA-approved targeted therapies effective against this BC subtype [5, 7]. Likewise, recurrent TNBCs frequently acquire resistance to standard-of-care therapeutic agents through mechanisms that remain incompletely understood. Recent studies have, however, shown that a subset of TNBC tumors that express active retinoblastoma gene (Rb1) can be sensitized to radiation therapy when treated with CDK4/6 inhibitors [8, 9].

The Y Box binding protein 1 (YB1) is a member of the highly conserved Cold Shock Domain protein family [10] with multifunctional properties both in the cytoplasm and inside the nucleus [10–18]. YB1 is also involved in various cellular functions, including regulation of transcription, mRNA stability, and splicing [10, 11, 15, 17] . Consistent with its essential biological functions, targeted disruption of *Yb1* in mice causes severe developmental defects and embryonic lethality[19]. Several studies have associated YB1 with the regulation of the malignant phenotype, cancer development, progression and metastasis, and drug resistance in several cancers, including ones of breast tissue origin [20–30]. More, importantly, an ongoing phase II NCI clinical trial (NCT02157051) is currently evaluating the efficacy of a YB1-based vaccine therapy targeting patients with advanced HER2-negative breast cancer, which further strengthens the significance of YB1 in BC biology and, therefore, the present studies. Our published studies showed YB1 to play a major role in driving the oncogenic behavior of TNBC tumors[30]. A study by Tailor et al., identified SU056 as a small molecule inhibitor that specifically binds to YB and lead to its degradation [31]. In that study, SU056 was shown to be effective in inhibiting the growth of ovarian cancer tumors both in vitro and in vivo[31]. A recent study has shown that SU056 can also inhibit breast cancer tumors [18]. In the present study we show that SU056, in combination with CDK4/6 inhibitor palbociclib, can be very effective in inhibiting tumor growth and metastasis of TNBC tumors, both in vitro and in pre-clinical models of TNBC tumors, including patient-derived xenografts (PDX) models. Mechanistically, we show that SU056 exerts its effect through the YB1-mediated regulation of expression genes involved in cell cycle progression, i.e., Cyclin D1 and CDK4, while palbociclib targets Rb1 for its degradation in Rb-positive TNBC cells, and PDXs.

Prospectively, these findings have important clinical implications, as they identify a novel, mechanism-based combination strategy that targets cell-cycle regulation in TNBC. The demonstrated efficacy of SU056 and palbociclib in clinically relevant PDX models supports the potential for patient stratification based on Rb1 status and provides a strong preclinical rationale for advancing this combination toward early-phase clinical trials in TNBC patients, particularly those with Rb-positive tumors and limited therapeutic options.

## Materials and Methods

### Cell lines and reagents

MDA-MB-231, MDA-MB-468 and 4T1 cells were procured from the American Type Culture Collection (ATCC; Manassas, VA) and maintained in accordance with the manufacturer’s specified protocols. Although cell line authentication was not explicitly conducted, we relied on the manufacturer’s quality control assurances. Periodic testing for Mycoplasma contamination was performed every 9 to 12 months. All cells were cultured at early passages (no more than 15), and each culture was passaged no more than five times before introducing a fresh vial. YB1-deficient cells were generated through electroporation of cancer cells with a ribonucleoprotein mixture of guide RNAs (sgRNA) and Cas9 (Synthego), following the manufacturer’s instructions [30]. A pool of three verified sgRNAs was used for each human or mouse gene (Synthego), with scrambled sgRNAs serving as a negative control. Western blot (WB) analysis validated efficient and stable knockout (KO). In cases where knockdown efficiency was below 80%, a second round of sgRNA delivery was implemented. No antibiotic selection was required, as the knockdown efficiency was sustained throughout the cells’ utilization. Primary antibodies for YB1 (Abcam, ab76149, 1:1000), CDK4 (Cell Signaling Technology, #12790 1:1000), Cyclin D1 (Cell Signaling Technology, #55506 1:1000), phosphor-Rb1 (Cell Signaling Technology, #8516 1:1000). Goat horseradish peroxidase-conjugated anti-mouse IgG and goat horseradish peroxidase-conjugated anti-rabbit IgG for western blot (Bio-Rad). Gel electrophoresis reagents for protein and DNA were from Bio-Rad. SU056 compound was synthesized in house in Dr. Su’s laboratory according to the formula published in [31]. 10mM stock solutions were prepared in DMSO and stored at -80C. Palbociclib was purchased from Selleckchem in powder form, dissolved in DMSO in stock solutions of 10mM, and stored aliquots at -80C. Working solutions were prepared as needed and stored in single use aliquots at -20C.

### MTT assay

Cell proliferation was assessed to determine the IC_50_ of SU056 and to evaluate the effects of SU056 combined with palbociclib for each cell line. MDA-MB-231, MDA-MB-468, and 4T1 (10,000 cells per well) were seeded into 96-well plates, treated the next day, and cultured for 48 hours. DMSO was used as a vehicle (negative control), and Cisplatin was used as a cytotoxic (positive) control. For combination treatments, SU056 was maintained at a fixed concentration of 1uM. Palbociclib was tested at varying concentrations from 0.5uM to 10uM for both the IC_50_ and the combination experiments. A stock solution of filter sterilized 5mg/mL MTT (3-(4,5-dimethylthiazol-2-yl)-2,5-diphenyl tetrazolium bromide) was prepared by dissolving 5mg MTT per 1mL phosphate-buffered saline (PBS). A working solution of MTT was prepared by mixing 25uL of MTT stock solution with 175uL of culture medium per well. The working solution was prepared in sufficient volume to treat all wells. After two hours of incubation with MTT, the supernatant was removed and DMSO was added to dissolve formazan crystals. Absorbance was measured at 570nm using a microplate spectrophotometer. All steps involving MTT, including subsequent incubation and solubilizing, were performed in the dark. All subsequent experiments used palbociclib at a concentration of 1uM.

### Colony formation assay

MDA-MB-231, MDA-MB-468, and 4T1 (15,000 cells) were seeded into 6-well plates, cultured for 7 days, fresh medium was supplemented every 3 days. Clones were washed with PBS, fixed with 4% paraformaldehyde (PFA) at room temperature for 20 minutes, and stained with 0.25% crystal violet solution for 20 minutes. Image acquisition of the 6-well plates were performed using the ChemiDoc MP Imaging system (Bio-Rad) and ImageJ software. Stained colonies were dissolved with 1% Sodium Dodecyl Sulfate (SDS) detergent and quantified using a microplate spectrophotometer at 590nm to compare absorbance relative to the control.

### Limiting dilution assay

Limiting-dilution assay (LDA) was performed as follows. Cells were seeded into 12 wells of a 96-well, ultra-low attachment plate at the following densities: 1024, 512, 256, 128, 64, 32, 16, or 8 cells/well for MDA-MB-231 and 350, 175, 87, 43, 21, 10, 5, or 2 cells/well for 4T1. The seeded cells were treated with SU056, palbociclib or both every 3 days and quantified on day 14 by light microscopy. Cancer stem cells frequencies were calculated using extreme limiting-dilution assay (ELDA) analysis software (http://bioinf.wehi.edu.au/software/elda/index.html).

### 3D-tumorsphere growth and invasion assays

For 3D single-tumorsphere formation, cells were seeded into a 96-well ultralow attachment (ULA) plate and monitored for 12 days, as described previously [32]. Invasion assays involved supplementing tumorsphere cultures with Matrigel, and invasion was monitored for 10 additional days. Images were captured and quantified using ImageJ software, as described previously [32].

### *In vivo* tumor growth and metastasis study

All in vivo experiments were approved by the Institutional Animal Care and Use Committee at the Cleveland Clinic Lerner Research Institute and Case Western Reserve University School of Medicine. NOD/scid/γ (NSG) female mice and BALB/C mice were purchased from The Jackson Laboratory, Bar Harbor, ME, USA and used for tumor growth and metastasis studies as described in our published studies [30, 33–35]. Mice were housed in microisolator units maintained on a 12 h light/dark cycle. Mice were given standard chow and water. For the cell xenograft models, parental and derivative cells were injected into mammary fat pads, in both the right and left sides (5 mice per group, 10 tumors per group) and treated as described below. Mice were sacrificed when to maximum tumor burden was reached in the control group (not to exceed 1500 mm^3^), and tumors and lung metastases were analyzed as described [30, 33–36]. For PDX models, tumor pieces from TM00098, purchased from The Jackson Laboratory, Bar Harbor, ME, USA, or AA0009, obtained from University Hospitals, Cleveland, OH, USA, were orthotopically xenografted in both inguinal mammary fat pads of adult female NSG mice. For treatment, mice with palpable tumors (200–300 mm3) were randomized into vehicle, palbociclib, SU056, or the combination of palbociclib and SU056 treated groups. Palbociclib was dissolved in 10% DMSO, 40% PEG300, 5% tween-80 and 45% saline, and delivered by oral gavage (p.o.) at 50 mg/kg, 5 days per week. SU056 was dissolved in 10% DMSO and 90% saline and injected intraperitoneally at 40 mg/kg, 5 days per week. Tumor size was measured by calipers twice weekly and volume was calculated as (length × width2)/2. To assess overt toxicity, mouse weight was measured once per week. At the experimental endpoint, each tumor was cut in half; one half was flash frozen and the second half was preserved in 4% paraformaldehyde (Thermo Fisher, Carlsbad, CA).

### Statistical analyses

Statistical analyses were performed using GraphPad Prism (version 8.0) and SPSS (version 21.0). All experiments were conducted in triplicate, and variables were expressed as mean ± SD. Student’s t-test was used, and significance was considered at p < 0.05. ANOVA was used when comparing more than two groups with continuous variables, and significance was considered at p < 0.05

## Results

### The small molecule inhibitor SU056 inhibits the YB1 oncogenic activities in TNBC cell lines in vitro

We previously published a study [30] establishing YB1 as a major driver of TNBC tumors progression and metastasis. A study by Tailor et al. [31], identified SU056 as a small molecule specific inhibitor of YB1. Here we sought to determine the effect of SU056 (Fig. 1A) on the oncogenic behavior of TNBC tumors through the inhibition of YB1 expression. Treatment of human MDA-MB-231 (Fig. 1B) , MDA-MB-468 (Fig. 1C) and mouse 4T1 (Fig. 1D) TNBC cell lines with increasing concentrations of SU056 established an IC50 of ∼0.5-1 µM. Therefore, all the subsequent in vitro assays used 1 µM of SU056. Western blot analyses showed a significant decrease of YB1 protein expression in all three cell lines when treated with 1 µM SU056 for 24 h. (Fig. 1E-G). In fact, the level of inhibition of YB1 was close to that achieved through CRISPR/Cas9-mediated knockout of YB1 (YB1-KO) (Sup. Fig 1A-C), therefore establishing the effectiveness of SU056 in targeting YB1. Several studies, including ours [30] showed that loss of YB1 expression attenuated the oncogenic behavior of TNBC cell lines, including the ability of YB1 to regulate cancer cell stemness [20, 30]. Treatment of TNBC cell lines with SU056 significantly inhibited 3D-tumorsphere growth of MDA-MB-231 (Fig. 2A) and 4T1 cells (Fig. 2B). Thus, supporting the inhibitory effect of SU056 on the stemness of these TNBC cell lines. This effect was further confirmed in the colony formation assay (Fig. 2C & 2D), and the limiting dilution assay (Fig. 2E and 2F). Treatment with SU056 resulted in a significant inhibition of these cells to form colonies as it resulted in 70% (Fig. 2B, MDA-MB-231) and 85% (Fig. 2C, 4T1) reduction in the number of colonies. Treatment with SU056 also resulted in a significant inhibition of their stemness, ranging in more than 2.5-fold reduction for MDA-MB-231 cells (Fig. 2E) to more than 3.5-fold reduction in 4T1 cells (Fig. 2F). Additional in vitro assays, such as transwell migration further confirmed the ability of SU056 to inhibit the in vitro oncogenic behavior of these TNBC cell lines (Sup Fig 2A & 2B).

**Figure 1.**
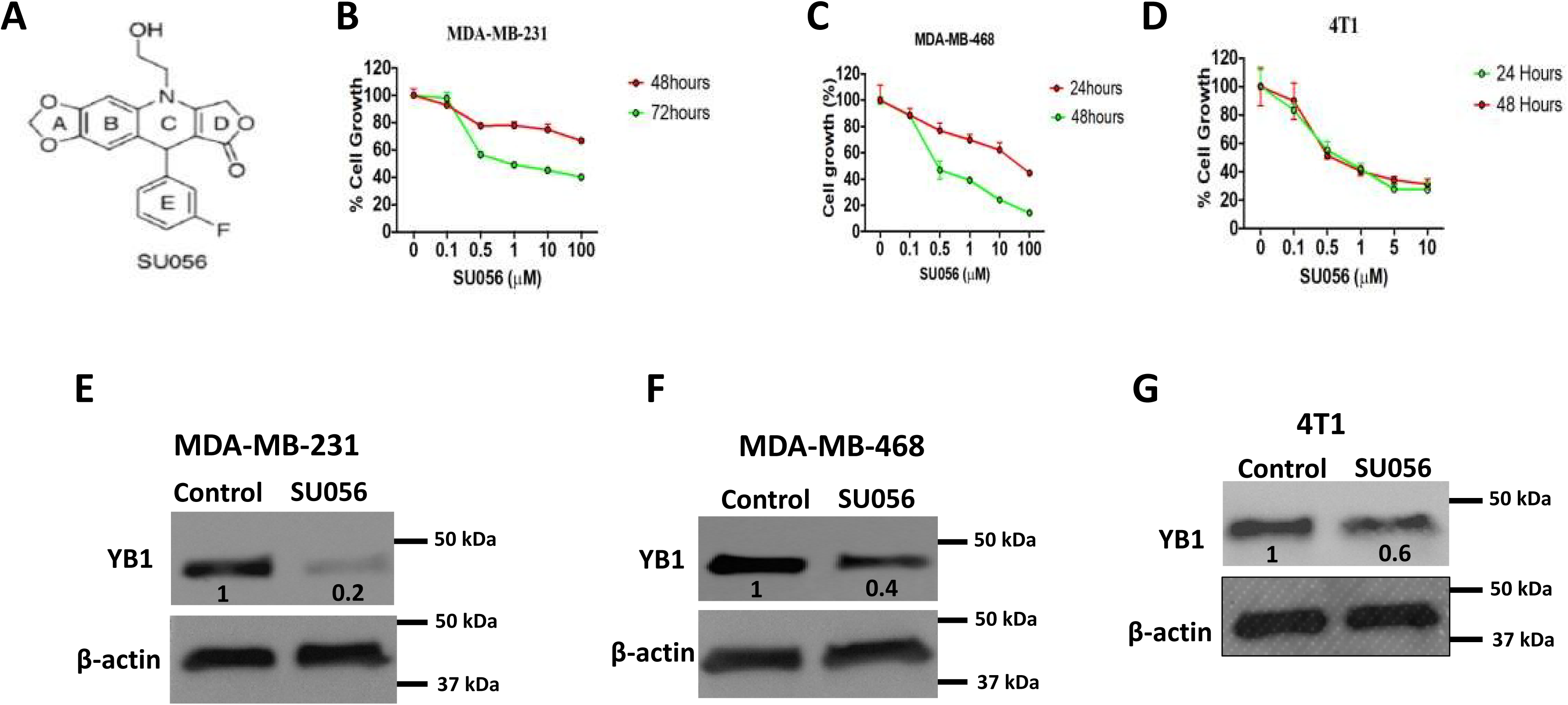
The small molecule inhibitor SU056 targets and inhibits YB1 protein in TNBC cell lines. **(A)** Chemical formula of SU056. **(B-D)** Quantification of cell proliferation using MTT assay of MDA-MB-231 (B), MDA-MB-468 **(C)** and 4T1 cells **(D)** under increased concentrations of SU056. Cell viability was plotted after 48 or 72 hours as a percentage of the cells seeded at day 0. Each datapoint is an average of at least 6 replicates. **(E-G)** Representative Western blots with anti-YB1 antibody of cell lysates of DMSO-treated (Control) or SU056-treated (1 uM) MDA-MDA-MB-231 **(E)**, MDA-MB-468 **(F)** or 4T1 cells **(G)** for 24 hours. β-Actin is a loading control. The numbers under each WB band represent the fold change in signal intensity with respect to its respective control band in each panel after normalization to the loading control signal. Data shown are representative of 3 independent experiments.

**Figure 2.**
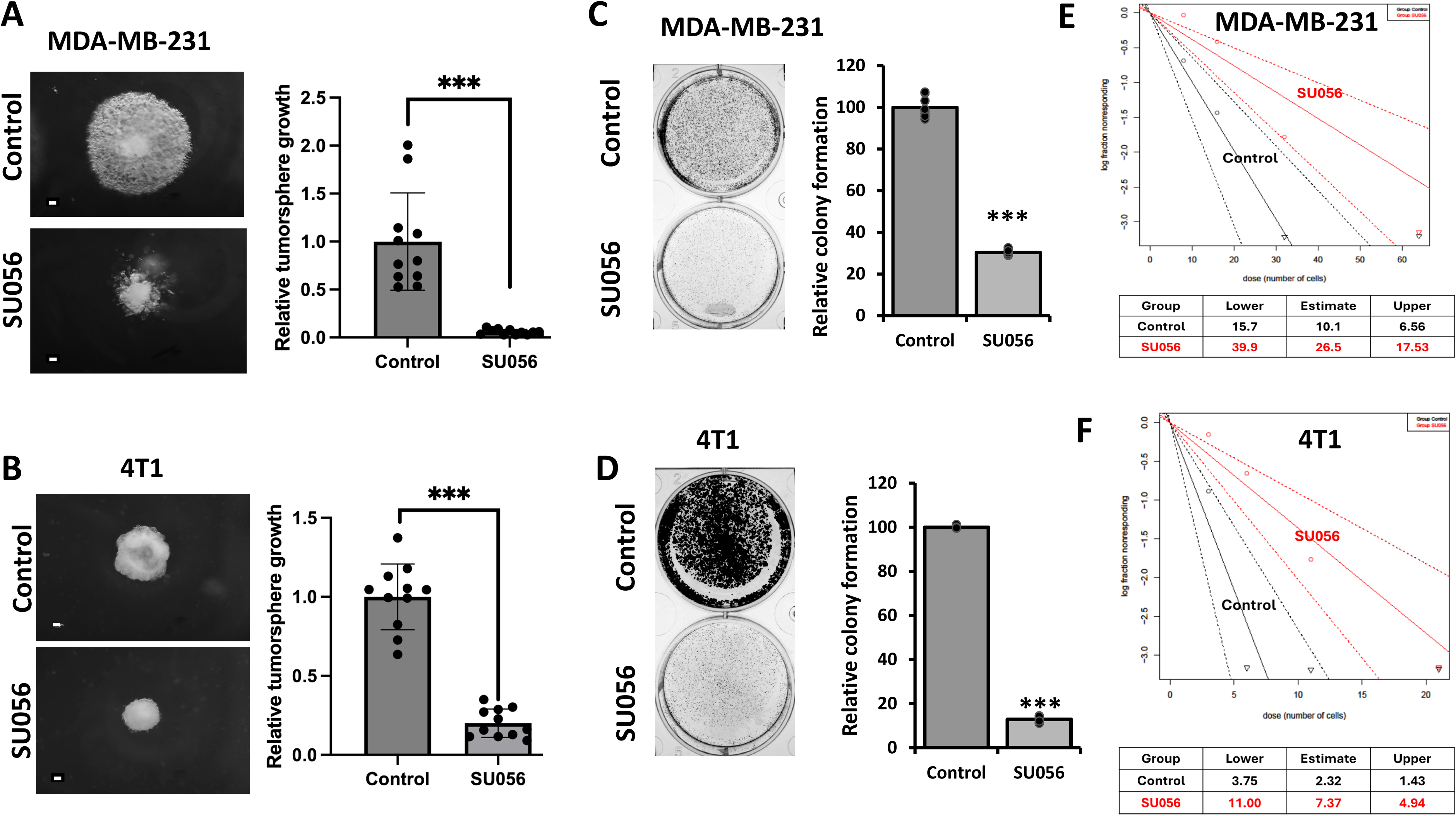
The small molecule inhibitor SU056 inhibits the YB1 oncogenic activities in TNBC cell lines in vitro. 3D-tumorsphere growth assay **(A-B),** 2D-colony formation assay **(C-D)**, and Limiting dilution assay **(E-F)** of Control MDA-MB-231 and 4T1 cells and their SU056-treated counterparts. Scale bar: 100 µm. Data are the mean ± SD (***p < 0.001, Student’s t test). Data shown are representative of 3 independent experiments.

### SU056 inhibits tumor growth and metastasis of TNBC tumors in vivo

To assess the clinical utility of SU056, we first established the in vivo toxicity parameters of SU056 in mice and found concentrations as high as 40 mg/kg can be tolerated by mice when injected intraperitoneally (IP), without noticeable toxicity or body wight loss. All our subsequent animal studies used a protocol of 10 mg/kg, IP injections twice a week for 4 weeks. NSG or BALB/c mice injected with MDA-MB-231 (Fig. 3A) or 4T1 cells (Fig. 3B), respectively showed a significant (p<0.1) decrease in tumor growth when treated with SU056, compared to mice treated with diluent DMSO. Metastasis to the lungs was also significantly (p<0.01) inhibited and both animal models (Fig. 3C-D). Mice injected with cells lacking YB1 expression (YB1-KO) also showed significant inhibition of tumor growth (Fig. 3A & 3B) and metastasis (Fig. 3C & 3D).

**Figure 3.**
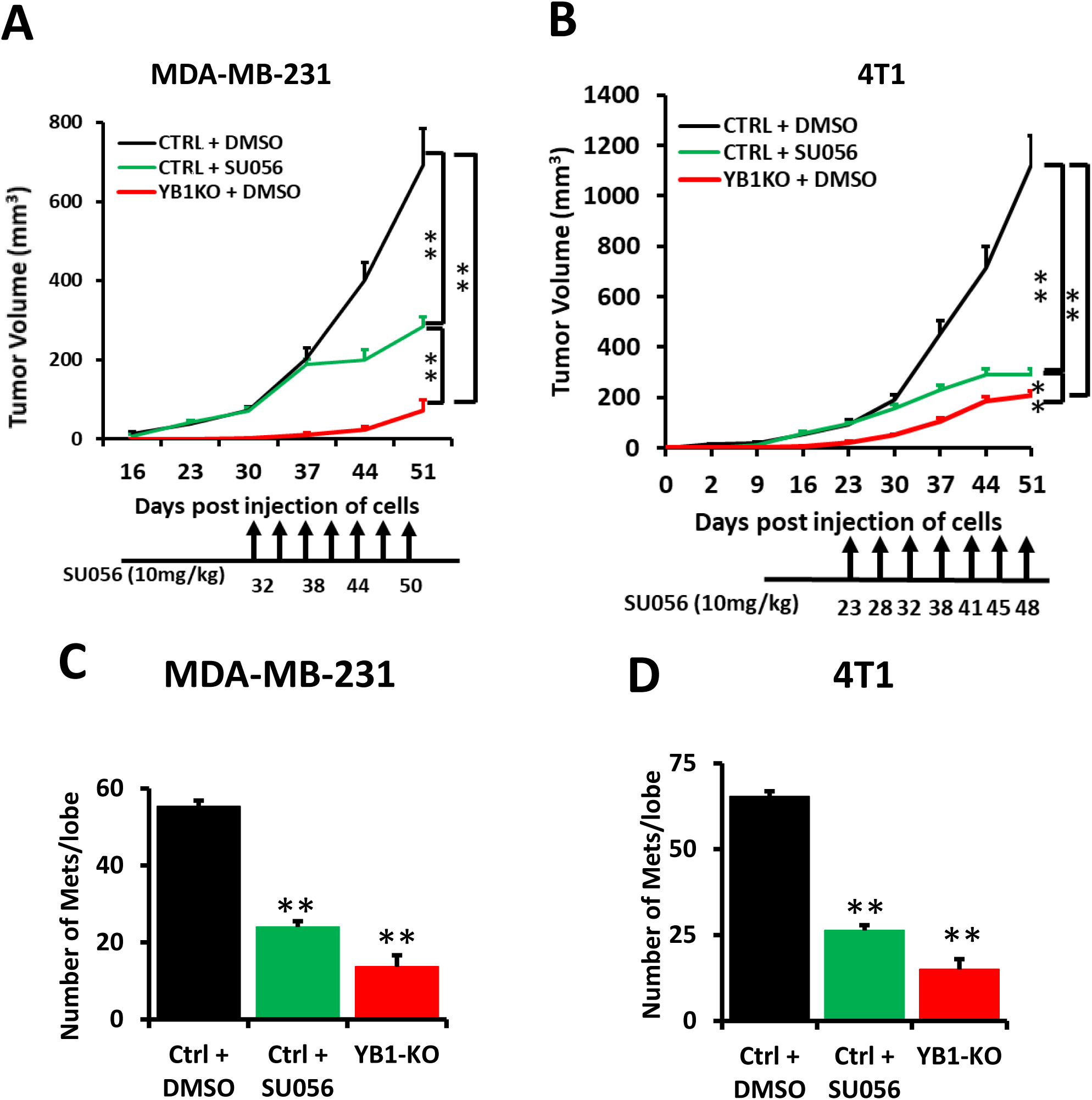
SU056 inhibits tumor growth and metastasis of TNBC tumors in vivo. (A-B) Tumor volume of NSG mice **(A)** or Balb/C mice **(B)** injected with control (Ctrl) MDA-MB-231 or 4T1 cells or their YB1-KO derivatives up to 50 days post injection. Once the average tumor volume reached ∼100 mm^3^ in the Ctrl mice, they were split into two groups, one group was injected (ip) with DMSO and the other group was treated with SU056 as indicated. The YB1-KO group was treated with DMSO. **(C-D**) quantification of metastatic foci of the lungs of each group from A and B, respectively. (**p < 0.01, ANOVA).

### SU056 Inhibits Cell-Cycle progression of TNBC Cells

Next, we tested the effect of SU056 on cell cycle progression. TNBC cells were treated with SU056 at concentrations of 1 μM for 24 hours, followed by cell cycle analysis. SU056 treatment resulted in a significant G0/1 phase arrest, evidenced by the accumulation of cells in the G0/1 phase. Both MDA-MB-231 (Fig. 4A) and 4T1 cells (Fig. 4B) showed increased G1/S phase arrest, with 75% of the cells accumulating in this phase after 24 hours of treatment compared to 50% in control cells. These results mimic the effect of YB1-KO (Sup. Fig. 3A & 3B). The effect of SU056 on cell cycle was attributable to the SU056-mediated downregulation of expression of cell cycle regulatory proteins Cyclin D1 and CDK4 (Fig. 4C and 4D). mRNA levels of CDK4 and Cyclin D1 were also inhibited after SU056 treatment in both MDA-MB-231 (Fig. 4E-F) and 4T1 cells (Fig. 4G-H). YB1-KO had similar effects (Sup Fig. 3C & 3D), suggesting that the YB1-mediated regulation of cell cycle genes is at the transcriptional level . Loss of expression of Cyclin D1 and CDK4 was also sustained in the tumors derived from mice injected with MDA-MB-231 (Fig. 4E) or 4T1 cells (Fig. 4F) and treated with SU056, similar to what was observed in the tumor derived from mice injected with YB1-KO MDA-MB-231 (Sup Fig. 3G) or YB1-KO 4T1 cells (Sup. Fig. 3H).

**Figure 4.**
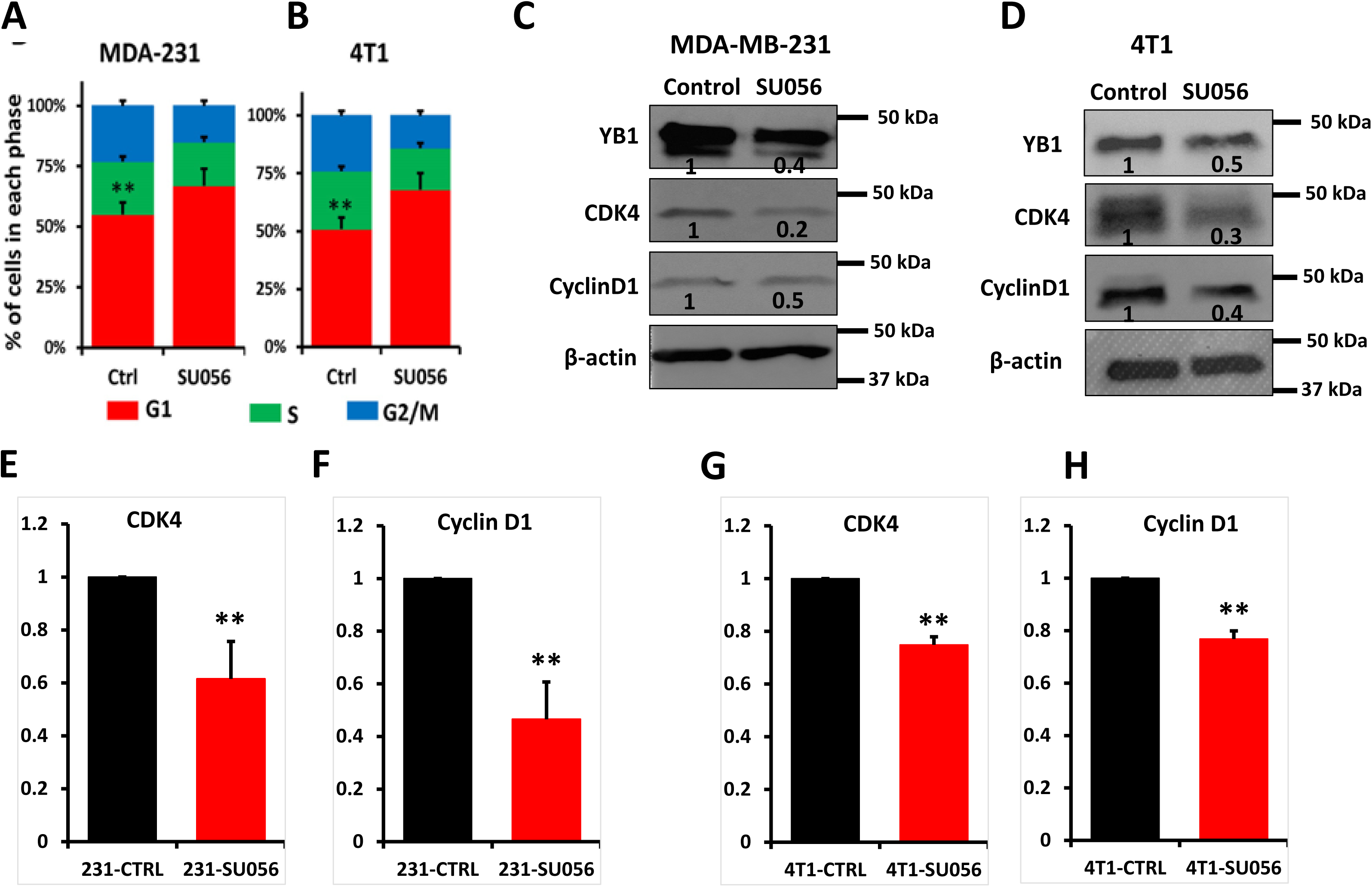
SU056 Inhibits Cell-Cycle progression and cell cycle genes in TNBC Cells. (A-B) Quantification of cells in each cell cycle phase of MDA-MB-231 **(A)** and 4T1 cells **(B)**. **(C-D)** Representative Western blots with the indicated antibodies of cell lysates of DMSO-treated (Control) or SU056-treated (1 µM) MDA-MDA-MB-231 **(C)** or 4T1 cells **(D)** for 24 hours. β-Actin is a loading control. The numbers under each WB band represent the fold change in signal intensity with respect to its respective control band in each panel after normalization to the loading control signal. **(E-H)** qt-RT-PCR of mRNA expression levels of CDK4 **(E and G)** and Cyclin D1 **(F and H)** from DMSO-treated MDA-MB-231 or 4T1 cells, or their SU056-treated counterparts. Data are the mean ± SD (**p < 0.01, Student’s t test). Data shown are representative of 3 independent experiments.

### Combination of SU056 with CDK4/6 inhibitor (Palbociclib) suppressed the growth through the Cyclin D1-CDK4/6 pathway in TNBC in Vitro

CDK4/6 inhibitors, such as palbociclib are potent inhibitors of cell cycle progression, and are routinely being used in the clinical setting for the treatment of patients with hormone-positive BC tumors [37–39]. However, recent small animal pre-clinical studies have established the efficacy of CDK4/6 inhibitors in the treatment of TNBC tumors in combination radiation therapy [8]. Given our data that shows SU056 to inhibit cell cycle, we sought to assess the effect of combining SU056 with palbociclib on the oncogenic behavior of TNBC tumors. Building on the finding that SU056 effectively inhibits TNBC growth, we hypothesized that combining SU056 with a palbociclib will further enhance the therapeutic impact against TNBC tumors. To test this hypothesis, we conducted in vitro assays to determine the impact of SU056 alone and in combination with a palbociclib on the oncogenic behavior of TNBC cells. Treatment of MDA-MB-231 (Fig. A-B) or 4T1 cells (Fig. C-D) with either SU056 or Palbociclib significantly inhibited 2D colony formation and 3D-tumorsphere growth. Combining SU056 with palbociclib further inhibited tumorsphere growth when compared to treatment with either SU056 or palbociclib alone. Next, we performed a limiting dilution assay and found that while treatment of MDA-MB-231 cells (Fig. 5E) with either SU056 or palbociclib resulted in >3-fold (26.45 vs 8.69) and >2-flod (19.39 vs 8.69) loss of stemness, respectively, combination of SU056 with Palbociclib resulted in >4-fold (32.78 vs 8.69) loss of stemness. Similar trends were observed with 4T1 cells (Fig. F). These findings support the enhanced inhibitory effect of combining SU056 with Palbociclib on the oncogenic behavior of TNBC cells.

**Figure 5.**
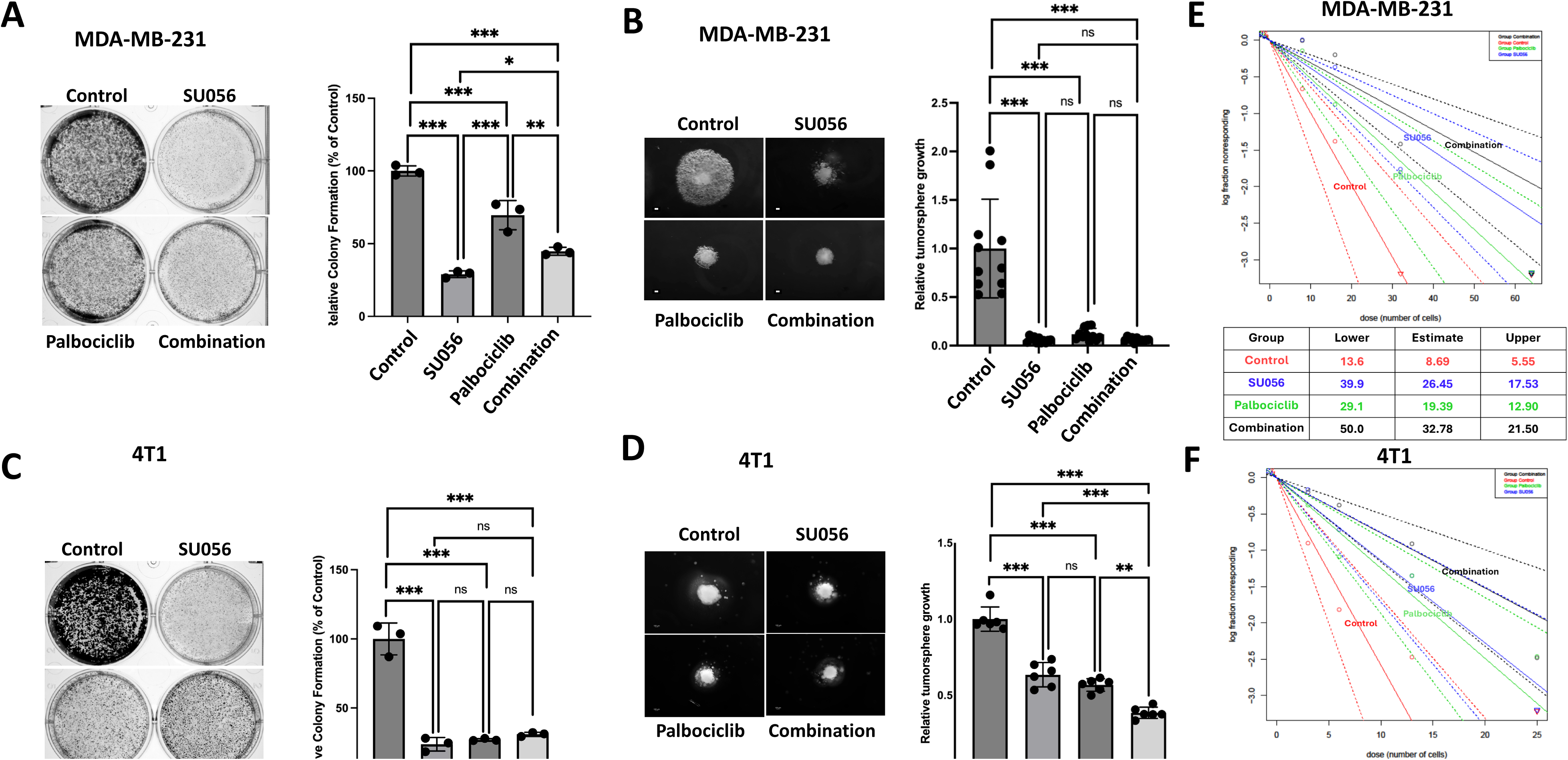
Combination of SU056 with CDK4/6 inhibitor (Palbociclib) suppressed the growth through the Cyclin D1-CDK4/6 pathway in TNBC in Vitro. 2D-colony formation assay **(A-B)**, 3D-tumorsphere growth assay **(C-D)**, and Limiting dilution assay **(E-F)** of Control MDA-MB-231 and 4T1 cells and their SU056-treated, palbociclib-treated, or SU056/palbociclib combination-treated counterparts. Scale bar: 100 µm. Data are the mean ± SD (*p < 0.05, **p < 0.01, ***p < 0.001, ANOVA). Data shown are representative of 3 independent experiments.

### Combination of SU056 with palbociclib suppressed growth and metastasis of TNBC tumors

The promising effectiveness of combined SU056 and palbociclib treatment in suppressing TNBC cells in vitro led us to evaluate the effect of this combination therapy on TNBC tumor progression in vivo. 4T1 TNBC cells were injected into the mammary fat pad area of female BALB/c mice. After allowing tumors to reach ∼100 mm^3^, mice were randomly assigned to one of four treatment groups (n=5 mice per group): vehicle control, SU056 alone, palbociclib alone, or a combination of SU056 and palbociclib. Treatments were administered as described in the Materials and Methods section.

In the vehicle-treated group, tumors continued to grow rapidly, and mice in this group were scarified 4 weeks post tumor cell injection, since they reach the tumor burden (>1200 mm^3^) allowed by the institutional animal protocol. While treatment with either SU056 or palbociclib alone showed a significant inhibition of tumor growth, mice treated with both of SU056 and palbociclib, tumor growth inhibition was further significant, compared to single treatment (Fig. 6A & 6B. At the end of the four-week treatment, tumors were collected and weighed to evaluate the impact of each treatment (Fig. 6C). While treatment with either SU056 or palbociclib alone resulted in ∼30 reduction of tumor weight, respectively, combination of SU056 with palbociclib resulted in >60% loss of tumor weight compared to the vehicle-treated mice. Together, these results support the use of SU056 and palbociclib combination therapy for the treatment of patients with TNBC tumors.

**Figure 6.**
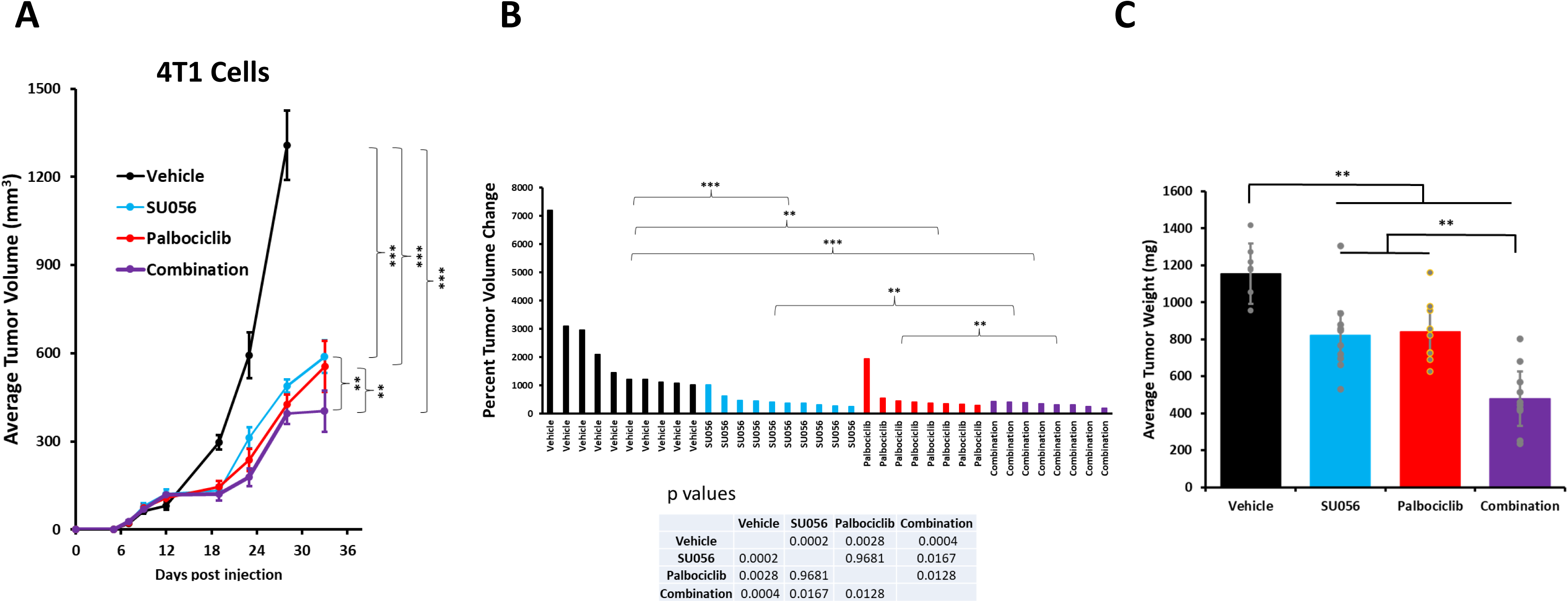
Combination of SU056 with palbociclib suppressed growth and metastasis of TNBC tumors. **(A)** ) Tumor volume of Balb/C mice injected with 4T1 in both left and right 4^th^ mammary gland. Once the average tumor volume reached ∼100 mm^3^ mice were split into 4 groups (n=5 mice per group), group 1 was treated with DMSO, group 2 was treated with SU056, group 3 was treated with palbociclib, and group 4 was treated with both SU056 and palbociclib. Drugs were administered via oral gavage as descried in the animal section of Materials and Methods. (**p < 0.01, ***p < 0.001, ANOVA). **(B)** Waterfall depicting the tumor volume change in each group at the end of experiment and plotted as the percent of the tumor volume just before the first treatment was administered. Statistical analyses are shown below. **(C**) Average tumor weight in each group at the end of experiment. (*p < 0.05, **p < 0.01, ANOVA).

### Combination of SU056 with palbociclib suppressed growth of TNBC PDX tumors

As a preliminary step towards the potential future use of SU056/palbociclib combination therapy in clinical trials, we assessed for its effectiveness on patient’s derived xenografts (PDX) in preclinical animal models. We used two PDX tumors: TM00098 and AA0009 that are maintained by Dr. Ruth Keri at the Cleveland Clinic. Of note, PDX TM00098 was derived from a tumor of a Caucasian American female patient, while PDX AA0009 was derived from a tumor of an African American female patient. For PDX TM00098, as shown by the relative tumor volume change (Fig. 7A), both single and combination treatments had significant inhibitory effect on tumor growth, with the combination therapy showing even more significant inhibitory effect than the single treatment. For PDX AA0009, however, neither single treatment, nor the combination treatment had any inhibitory effect on tumor growth (Fig. 7B). Further investigation for this discrepancy between the two tumor models, established that PDX TM00098 has an active Rb1 function, as shown by loss of Rb1 phosphorylation (Fig. 7C) similar to MDA-MB-231 (Fig. 7D) and 4T1 cells (Fig. 7E), when treated with palbociclib, while PDX AA0009 (Fig. 7C) and MDA-MB-468 (Fig. 7E) did not show any sign of Rb1 activity. Given that palbociclib along with other CDK4/6 inhibitors specifically target Rb1 (Fig. 8), the absence of Rb1 in PDX AA0009 negated the effect of palbociclib, see below for further discussion.

**Figure 7.**
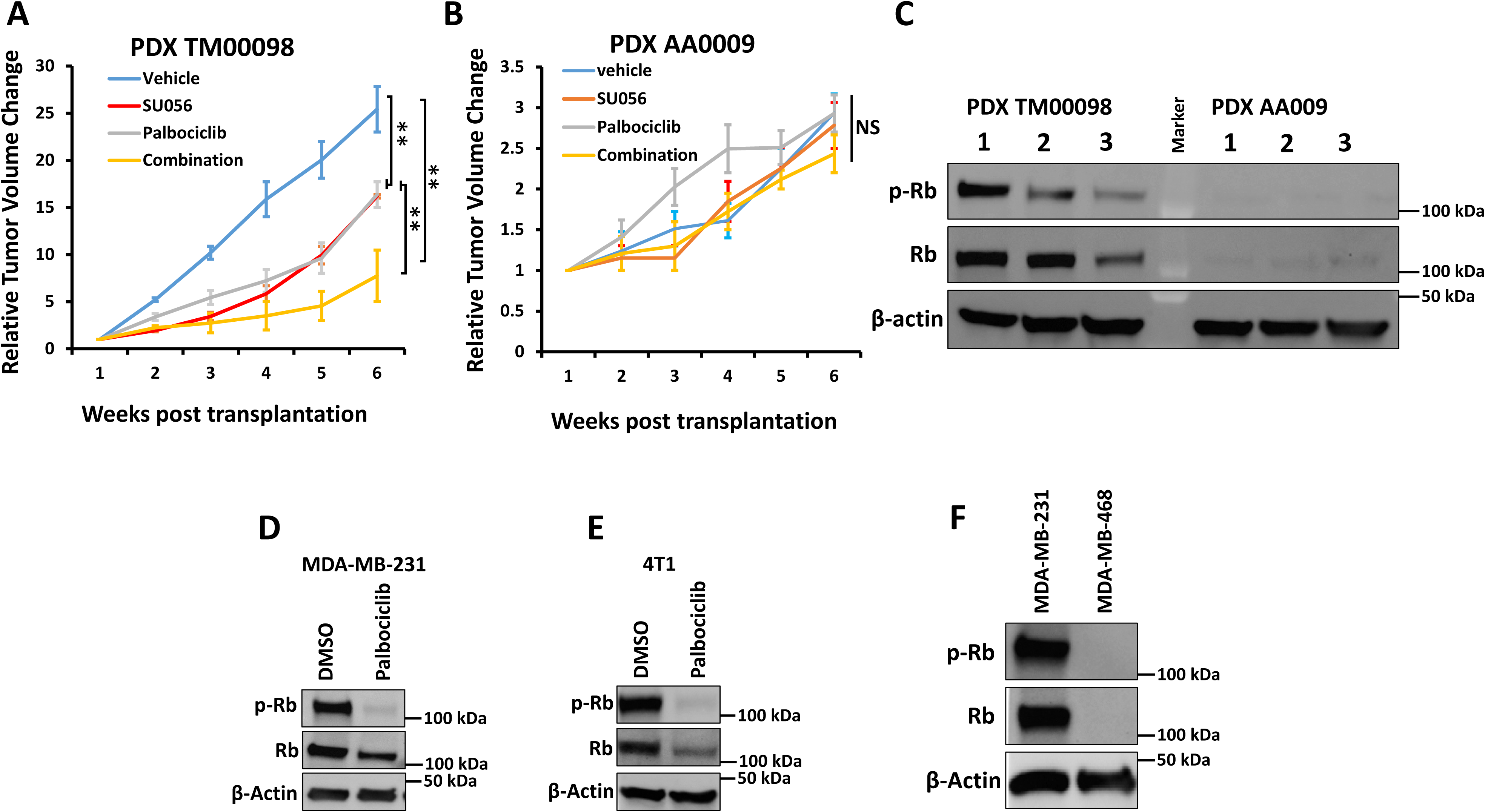
Combination of SU056 with palbociclib suppressed growth of TNBC PDX tumors. Tumor volume of NSG mice injected with PDX TM00098 (A) or PDX AA0009 in 4th mammary gland. Once the average tumor volume reached ∼100 mm^3^ mice were split into 4 groups (n=10 mice per group), group 1 was treated with DMSO, group 2 was treated with SU056, group 3 was treated with palbociclib, and group 4 was treated with both SU056 and palbociclib. Drugs were administered via oral gavage as descried in the animal section of Materials and Methods. (NS, not significant; **p < 0.01, ANOVA). **(C)** Representative Western blots with anti-Rb or anti-pRb antibodies of cell lysates from naïve TM00098 or AA0009 tumors. **(D-E)** Representative Western blots with anti-Rb or anti-pRb antibodies of cell lysates of DMSO-treated (Control) or palbociclib-treated MDA-MDA-MB-231 **(D)** or 4T1 cells **(E)** for 24 hours. (F) Representative Western blots with anti-Rb or anti-pRb antibodies of cell lysates of MDA-MB-231 or MDA-MB-468 cells. β-Actin is a loading control.

**Figure 8.**
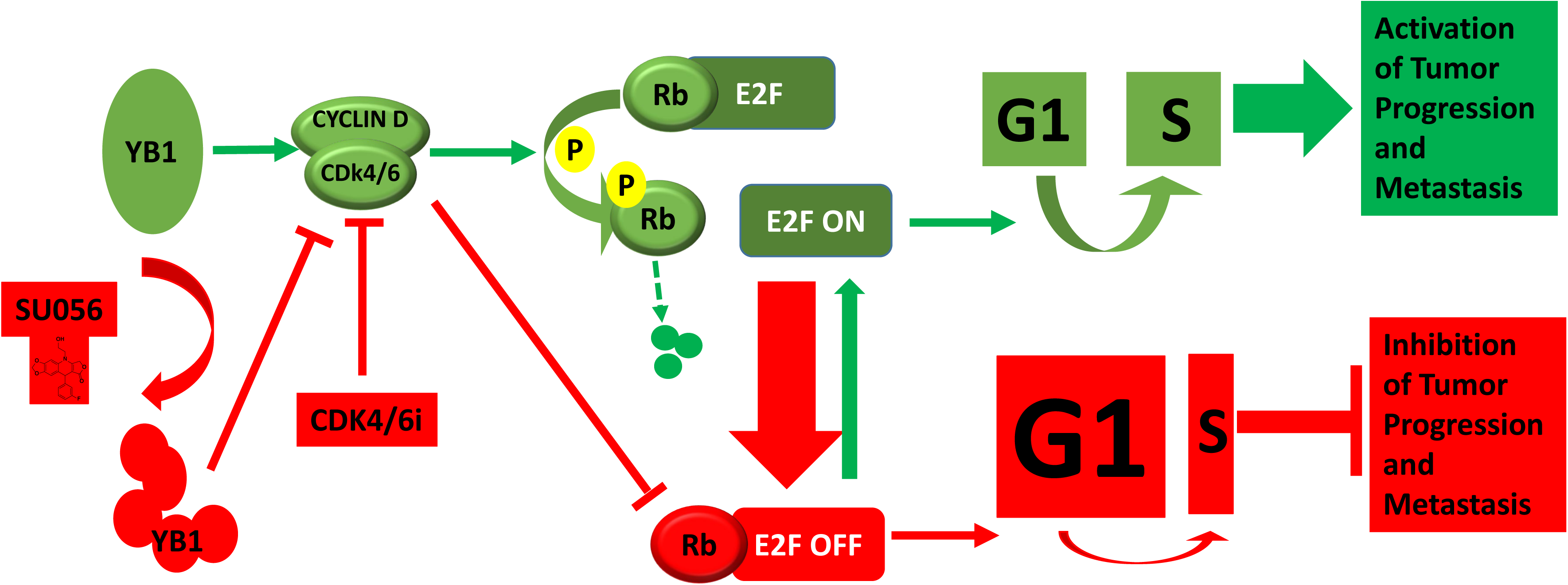

## Discussion

The absence of validated molecular targets in TNBC has prompted intense efforts to elucidate the key pathways that drive tumor growth, survival, and therapeutic resistance. Identifying such pathways is critical for developing rational, mechanism-based treatment strategies capable of improving patient outcomes. One promising target that has emerged from these efforts is Y-box binding protein-1 (YB1), a multifunctional transcription and translation factor implicated in oncogenic signaling, cell-cycle progression, stress adaptation, and chemoresistance across multiple cancer types, including breast cancer. Elevated YB1 expression has been associated with aggressive tumor behavior, poor prognosis, and resistance to conventional therapies, positioning YB-1 as an attractive therapeutic target in TNBC.

SU056 is a small-molecule inhibitor that targets YB1 and has demonstrated potent anti-tumor activity in several cancer models [18, 31]. In the present study, we investigated the therapeutic potential of SU056 in TNBC, hypothesizing that pharmacologic inhibition of YB1 would disrupt oncogenic behavior of TNBC tumors and impair their survival. Consistent with this hypothesis, we found that low concentrations of SU056 (<1 µM) effectively suppressed the levels of YB1 protein in several TNBC cell lines to levels comparable to the genetic targeting of YB1 (CRISPR-mediated knockout). The effect of SU056-mediated degradation of YB1 was reflected in the significant inhibition of the oncogenic behavior TNBC cells, using a combination of 2D colony formation, trans-well migration and Matrigel invasion assays, and 3D tumorsphere growth and invasion assays. The magnitude of inhibition of these oncogenic activities by SU056 was comparable to that achieved by CRISPR-mediated genetic knockout o YB1, further supporting the effectiveness and specificity of SU056.

Mechanistically, we established that SU056-mediated targeting of YB1 induced a robust G0/G1 cell-cycle arrest in TNBC cell lines, through the inhibition of expression of cell cycle gens Cyclin D1 and CDK4, which indicates that YB-1 inhibition impairs key functional properties associated with tumor aggressiveness and metastatic potential. Importantly, the anti-tumor activity of SU056 extended beyond in vitro systems. In vivo studies using mouse models of TNBC demonstrated that SU056 significantly restrained tumor growth, providing strong evidence for its therapeutic efficacy and validating YB-1 as a druggable vulnerability in TNBC.

CDK4/6 inhibitors, which induce G0/G1 cell-cycle arrest by blocking cyclin-dependent kinase activity, have shown remarkable success in hormone receptor–positive breast cancer but limited efficacy as monotherapy in TNBC. Based on published studies showing a potential effect CDK4/6 inhibitors on Rb^+^-TNBC tumors [8], and based on our on the data showing that SU056 also targets cell cycle, we posited that simultaneous inhibition of YB1 and cell cycle progression may lead to enhanced inhibition of the oncogenic activities of Rb1^+^ TNBC cell lines and tumors. Indeed, the combination of SU056 and palbociclib achieved those results both at the cell line and the xenograft levels. Both MDA-MB-231 and 4T1 express wildtype Rb1, and were responsiveness to palbociclib where a significant decrease of pRb1 levels was observed after treatment with palbociclib (Fig. 7D). MDA-MB-468, which does not express Rb1 (Fig. 7F), was not responsive to palbociclib not shown). Similarly, in our PDX models, only PDX TM00098 which expressed Rb1 (Fig. 7C) showed a significant inhibition in tumor growth in animals treated with palbociclib (Fig. 7A). More importantly, the combination of SU056 with palbociclib resulted in tumor growth inhibition that was more significant than the treatment with either SU056 or palbociclib alone. Although we were not able to determine synergy, the results are very promising. Additional experiments are needed to determine the optimal dosing conditions and treatment modalities. On the other hand, in the PDX model AA0009 which does not express Rb, treatment with palbociclib did not have any effect on tumor growth.

One final observation to point to is that MDA-MB-231 and PDX TM00098 are derived from Caucasian American patients, while MDA-MB-468 and PDX AA0009 were derived from African American patients. Both MDA-MB-231 and PDX TM00098 express Rb1 and are responsive to CDK4/6 inhibitors, while both MDA-MB-468 and PDX AA0009 do not express Rb1 and did not respond to CDK4/6 inhibitors. While this observation is based on an n of 2, one may suggest that screening for Rb negativity may be more prevalent in African American women with TNBC. Therefore, genetic screening for Rb1 expression in TNBC patients may identify those who may benefit from treatment CDK4/6 inhibitors, especially in African American women.

Collectively, our findings identify combined inhibition of YB1 and CDK4/6i as a compelling therapeutic strategy for TNBC. The robust activity observed in clinically relevant preclinical models provides a strong rationale for further translational development and supports the evaluation of YB1–CDK4/6i combination therapy in future clinical studies aimed at improving outcomes for patients with TNBC. Importantly, the dependence of therapeutic efficacy on Rb1 status suggests a biomarker-driven strategy for patient selection, enabling identification of TNBC patients most likely to benefit from SU056–CDK4/6i therapy.

## List of Abbreviations

BC: Breast cancer
TNBC: Triple negative breast cancer
ER: Estrogen receptor
PR: Progesterone receptor
HER2: Human epidermal growth factor receptor 2
YB1: Y-box binding protein-1
PDX: Patient-derived xenograft
CDK4: Cyclin-dependent kinase 4
2D: 2-Dimensional
3D: 3-Dimensional
CRISPR: Clustered Regularly Interspaced Short Palindromic Repeats
KO: Knockout
Rb1: Retinoblastoma gene 1.
Ab: Antibody
sgRNA: small guide RNA
mRNA: Messenger RNA
qRT-PCR: quantitative reverse transcription polymerase chain reaction

## Declarations

### Ethical approval

All animal studies were performed under protocols approved by the Institutional Animal Care and Use Committee at Case Western Reserve University and MetroHealth System.

### Author’s contributions

WW: Performed experiments, acquired data, analyzed data, co-wrote the first draft of the manuscript; YK: Performed experiments, acquired data, analyzed data; KMO: Performed experiments; KWB: Performed experiments; BS: Designed the research studies and provided reagents; RK: Designed the research studies and provided reagents; KS-A: Designed the research studies, performed experiments, acquired data, analyzed data, provided reagents, and wrote the final version of the manuscript. All authors reviewed and approved the manuscript.

## Supporting information

Supplementary Figures with Legends

## Acknowledgements

We thank Hana Arabi, Justin Szpendyk and Cody M. Orahoske for their technical assistance. We also thank members of K. Sossey-Alaoui lab for their critical inputs and. This work was supported in part by grant R01CA226921, grant R01CA272621, METAvivor Translational Research Award, and MetroHealth startup funds to K. Sossey-Alaoui.

## Availability of data and materials

All data are contained within the article. Requests for reagents should be addressed to K Sossey-Alaoui.

## Supplementary figures legends

**Sup. Fig. 1:** Representative Western blots with anti-YB antibodies of cell lysates from parental MDA-B-231 (A), 4T1 (B) and MDA-MB-468 (C) cells, their YB1-KO derivatives or parental cells treated with SU056 (1uM) for 24 hours). β-Actin is a loading control.

**Sup. Fig. 1: Representative images of 3D transwell migration assays of MDA-MB231 (A) or 4T1 cell (B)** , their YB1-KO derivatives or parental cells treated with SU056 (1uM) for 24 hours). Quantification of migrated cells in each condition is also shown. Data are the mean ± SD (***p < 0.001, ANOVA). Data shown are representative of 3 independent experiments.

Sup. Fig. 3: Quantification of cells in each cell cycle phase of parental MDA-MB-231 **(A)** and 4T1 cells **(B)**, or their YB1-KO derivatives. **(C-D)** Representative Western blots with the indicated antibodies of cell lysates of parental MDA-MB-231 **(C)** and 4T1 cells **(D), or their YB1-KO derivatives**. β-Actin is a loading control. (E-F) Representative Western blots with the indicated antibodies of cell lysates of tumors derived from mice injected with parental MDA-MB-231 **(E)** and 4T1 cells **(F)**, or their YB1-KO derivatives. β-Actin is a loading control. Representative Western blots with the indicated antibodies of cell lysates of tumors derived from mice injected with parental MDA-MB-231 **(E)** and 4T1 cells **(F)** and treated with DMSO, or their SU056-treated counterparts. β-Actin is a loading control.

