## Supplementary Figures with Legends for "The small molecule inhibitor SU056-mediated targeting of YB1 inhibits the Rb pathway in triple negative breast cancer tumors"

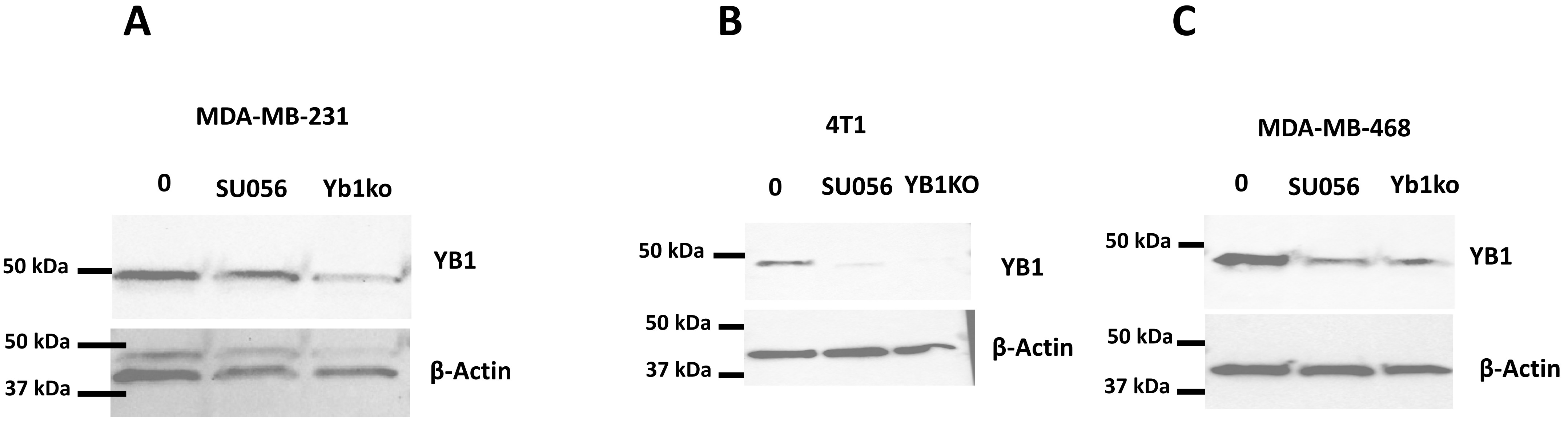
A
MDA-MB-231
0
SU056
Yb1ko
50 kDa
YB1
50 kDa
37 kDa
β-Actin
B
4T1
0
SU056
YB1KO
50 kDa
YB1
50 kDa
37 kDa
β-Actin
C
MDA-MB-468
0
SU056
Yb1ko
50 kDa
YB1
50 kDa
37 kDa
β-Actin

Sup. Fig. 1

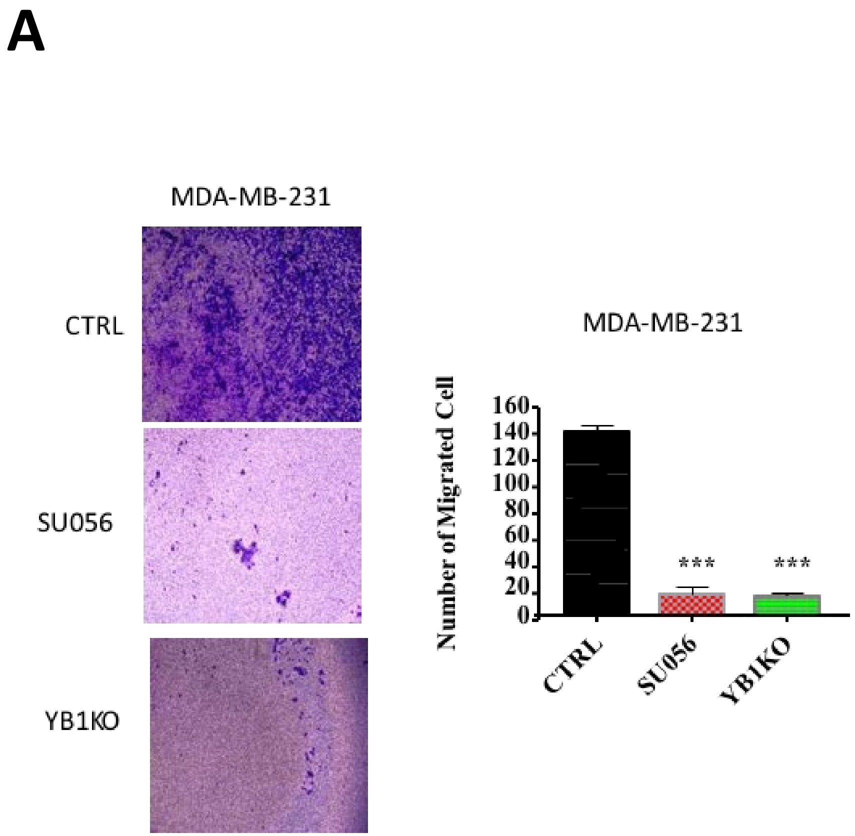
A
MDA-MB-231
CTRL
SU056
YB1KO
MDA-MB-231
Number of Migrated Cell
160
140
120
100
80
60
40
20
0
***
***
CTRL
SU056
YB1KO

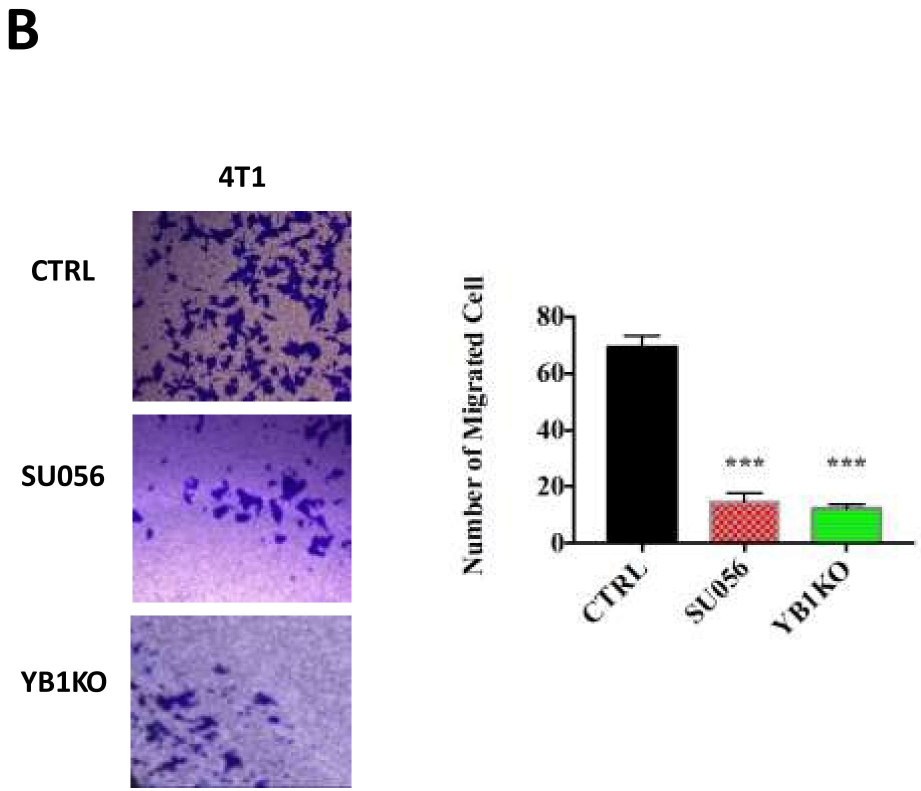
B
4T1
CTRL
SU056
YB1KO
Number of Migrated Cell
80
60
40
20
0
***
***
CTRL
SU056
YB1KO

Sup Fig. 2

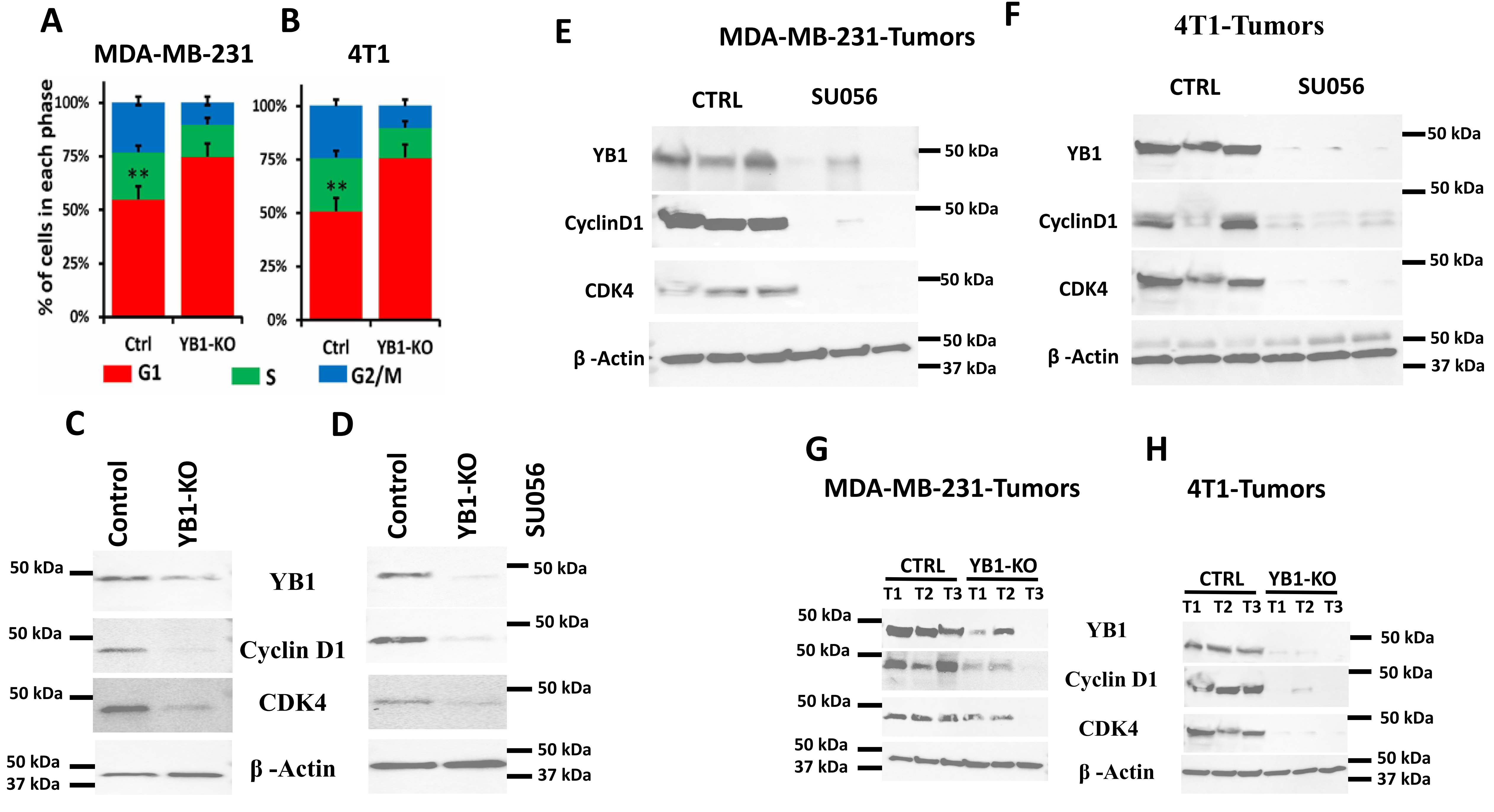

A
MDA-MB-231
B
4T1
% of cells in each phase
100%
75%
50%
25%
0%
**
Ctrl
YB1-KO
G1
S
G2/M
E
MDA-MB-231-Tumors
CTRL
SU056
YB1
CyclinD1
CDK4
β -Actin
50 kDa
37 kDa
F
4T1-Tumors
C
Control
YB1-KO
Cyclin D1
MDA-MB-231
D
SU056
4T1
G
MDA-MB-231-Tumors
T1
T2
T3
H
4T1-Tumors


Sup. Fig. 3

**Supplementary figures legends:**

**Sup. Fig. 1:** Representative Western blots with anti-YB antibodies of cell lysates from parental MDA-B-231 (A), 4T1 (B) and MDA-MB-468 (C) cells, their YB1-KO derivatives or parental cells treated with SU056 (1uM) for 24 hours). β-Actin is a loading control.

**Sup. Fig. 2: Representative images of 3D transwell migration assays of MDA-MB231 (A) or 4T1 cell (B)** , their YB1-KO derivatives or parental cells treated with SU056 (1uM) for 24 hours). Quantification of migrated cells in each condition is also shown. Data are the mean ± SD (***$p < 0.001$, ANOVA). Data shown are representative of 3 independent experiments.

**Sup. Fig. 3:** Quantification of cells in each cell cycle phase of parental MDA-MB-231 **(A)** and 4T1 cells **(B)**, or their YB1-KO derivatives. **(C-D)** Representative Western blots with the indicated antibodies of cell lysates of parental MDA-MB-231 **(C)** and 4T1 cells **(D), or their YB1-KO derivatives**. β-Actin is a loading control. (E-F) Representative Western blots with the indicated antibodies of cell lysates of tumors derived from mice injected with parental MDA-MB-231 **(E)** and 4T1 cells **(F)**, or their YB1-KO derivatives. β-Actin is a loading control. Representative Western blots with the indicated antibodies of cell lysates of tumors derived from mice injected with parental MDA-MB-231 **(E)** and 4T1 cells **(F)** and treated with DMSO, or their SU056-treated counterparts. β-Actin is a loading control.
